# Reducing Offsite Modification using 2-mercaptoethanol for Proteome Analysis

**DOI:** 10.1101/2024.08.09.607156

**Authors:** Arisa Suto, Takashi Matsui, Yoshio Kodera

**Author notes:** Correspondence: Dr. Takashi Matsui, Postal address: 1-15-1 Kitasato, Minami-ku, Sagamihara, Kanagawa 252-0373, Japan.

## Abstract

Alkylation of the thiol group in cysteine (Cys) residues using halide reagents is a significant step in proteomics. However, non-specific modifications to the N-terminus and other amino acids are known. Thus promiscuous offsite alkylation in the peptide further complicated the MS spectra and thus made the difficulty of identification and quantification of all peptides. 2-mercaptoethanol (2-ME) is not only a regent for the reduction of the disulfide bond but also is bound to the Cys residue. Furthermore, it is known that dimethyl sulfoxide (DMSO) enhances the disulfide bond formation. Thus, based on these facts, we developed a method for specifical modification of Cys residues using 2-ME and DMSO. The specific modification of Cys residue by 2-ME were promoted by the concentration-dependent manner of DMSO with quite less offsite modification reaction compared with recent procedures. This Cys-specific modification technique may not only improve the quantification of peptides containing cysteine but also enhance the quantification accuracy of all peptides.

## Introduction

The homeostasis in living organisms is maintained by the expression, degradation and post-translational modification of proteins. Therefore, analyzing the alternations in the quantity and quality of proteins is important to understand biological systems and the mechanisms of disease manifestation^1^. In quantitative proteomics, a method for analysis of the abundance of proteins in the target sample, the object and the reference samples are labeled by labeling reagents of different isotopic masses, respectively. This allows for analyzing changes in the abundance and modification states of proteins and peptides between healthy and diseased individuals using Liquid Chromatography-Mass Spectrometry (LC-MS), enabling the exploration of potential biomarkers and indicators of drug efficacy^2^. Recently, data-independent acquisition (DIA), a new LC-MS technique, has been developed to perform comparative analysis with greater sensitivity and depth on low-abundance proteins and peptides that had not been compared previously^3^. Other methods were developed to evaluate the protein conformation in solution by applying the quantitative proteomics^4,5^. Thus, quantitative proteomics is a fundamental technique that finds applications in various fields.

Although cysteine (Cys) residue contributes to the stability of the protein structure through disulfide bonds, it undergoes various modifications *in vivo* and *in vitro*^6^. To avoid various non-physiological modifications, disulfide bonds are cleaved and thiol groups are protected using alkylating reagents before LC-MS analysis. Although the halide compounds, such as 2-iodoacetamide (IAA), is commonly used in proteomics, it often cause side reactions to the N-terminus and other amino acids^7^. Furthermore, IAA unintentionally alkylates methionine (Met) under lower pH conditions^8^, suggesting the promiscuous offsite alkylation in the peptide further complicated the MS spectra and thus made the difficulty of identification and quantification of all peptides. Therefore, specific modification for the Cys residue was needed for a more accurate quantitative analysis of proteomics.

In this study, Utilizing the fact that 2-mercaptoethanol (2-ME) adds a +76 Da modification to the free thiol groups of Cys ^9,10^, and that dimethyl sulfoxide (DMSO) promotes the formation of disulfide bonds between Cys residues ^11^, we attempted to specifically induce the formation of disulfide bonds between the thiol groups of Cys and 2-ME under the presence of DMSO to prevent the scrambling modifications at Cys and other amino acids, and thus we achieved the development of a method for identifying a larger number of peptides coupled with performing a more quantitative LC-MS analysis using β-Galactosidases (β-Gal) as an example than present procedures.

## Materials and Methods

### Cysteine modification of β-Gal

5 µM of β-Gal (Sigma-Adrich, Missouri, USA) dissolved in 50 mM Tris-HCl, pH 8.0 was incubated for 0.5 h at 50℃. For the assessment of the effect of DMSO, 2.5 µM of β-Gal was incubated with 0.5 mM 2-ME and varying concentrations of DMSO (0-30%, Thermo Fisher Scientific, Massachusetts, USA) at room temperature for 1 h. For the efficiency of 2-ME adduction to the Cys residue, 2.5 µM of β-Gal was incubated with 2-ME (0-4 mM) and 20% DMSO at room temperature for 1 hour. For the carbamidomethylation of Cys residue, 2.8 µM of β-Gal was alkylated with 20 mM IAA (Nacalai Tesque, Kyoto, Japan) and 20 mM triethylammonium bicarbonate (Thermo Fisher Scientific) for 1 h at room temperature in the dark.

### Preparation for LC-MS analysis

After digestion using 3.3 ng/µL chymotrypsin (Promega, Wisconsin, USA) and 5 mM CaCl2 at 25℃ overnight, the sample was mixed with a 1.5 × volume of 1.7% trifluoroacetic acid (TFA) and 1 pmol of synthetic peptide, YVYVADVAAK, as an internal standard. The mixture was desalted and eluted by 70% acetonitrile (ACN) and 0.1% TFA acid and subsequently freeze-dried., as the procedure described previously ^12^.

### LC-MS/MS analysis

After resolving the freeze-dried sample using the solvent (3% ACN and 0.1% Formic acid (FA)), 1 pmol of the resuspended sample was analyzed by a quadrupole Orbitrap benchtop mass spectrometer (Q-Exactive, Thermo Fisher Scientific) connected with an EASY-nLC 1000 system (Thermo Fisher Scientific) as followed the previously described^13^ with slight modifications. Briefly, peptides were eluted with a gradient of solvents A (0.1% FA)) and B (90% ACN and 0.1% FA): 0–1 min, 5–10% B; 1–20 min, 10–25% B; 20–26 min, 25–50% B; 26–27 min 50–90% B on an analytical column (C18, particle diameter 3 µm, 0.075 mm × 125 mm; Nikkyo Technos, Tokyo, Japan) at flow rate of 300 nL/min. MS1 spectra were collected over the scan range 350–1200 *m*/*z* at 70,000 resolution to hit an automatic gain control target of 1 × 10^5^. The 10 most intense ions with charge states of 2^+^ to 4^+^ and an intensity > 2×10^4^. MS2 spectra were acquired on the Orbitrap mass analyzer with a mass resolution of 35,000. A dynamic exclusion with a time window of 5 s was applied.

The raw data obtained from mass spectrometry have been deposited in the ProteomeXchange Consortium (http://proteomecentral.proteomexchange.org) via the jPOST partner repository (http://jpostdb.org)^14^ as set identifiers PXD054531for ProteomeXchange and JPST003251 for jPOST.

### Peptide identification and quantification

LC-MS/MS data were interrogated against the UniProt sequence database (release 31st July 2019, entry 557,016, all species, reviewed) using *de novo* sequencing algorithm incorporated into the PEAKS Studio version X (http://www.bioinfor.com/peaks-studio/, Bioinformatics Solutions, Waterloo, Canada)^15^ as the search engine. The parameters were followed: precursor mass tolerance, 6 ppm; fragment mass tolerance, 0.02 Da; enzyme, none; maximum missed cleavage sites, 3; variable modifications, methionine oxidation, carbamidomethylation (+57.02, IAA), 2-mercaptoethanol adduction (+76.00, 2-ME). The peptide identification was filtered to a false discovery rate (FDR) <1%.

Extracted ion chromatograms (XICs) for precursor ions were obtained using Skyline 22.2.0.527 (http://proteome.gs.washington.edu/software/skyline)^16,17^ based on the identified peptide library. The spectrum library was imported from the mzxml file generated by PEAKS with a cutoff score of FDR = 0.99. Peptide settings were followed: enzyme, Chymotrypsin FWYL/P; maximum missed cleavages, 9; minimal length of peptide, 4; maximal length, 25; modifications, oxidation (Met), 2-mercaptoethanol adduction (2-ME); maximum variable mods, 3. Transition settings were followed: precursor charges, 2^+^ to 4^+^; type, p (precursor); ion mass tolerance, 0.02 m/z; isotope peaks included, count 3; mass analyzer, Orbitrap, resolution, 70,000 at 200 m/z; use only scans within 5 min of predicted retention time; isotope labeling enrichment, default.

For the identification of the post-translational modification by 2-ME or IAA, the static modification of carbamidomethylation (+57.02, IAA), or 2-mercaptoethanol adduction (+76.00, 2-ME) was also applied to all amino acids and N- and C-terminus individually. The modified residue in a peptide was assigned from the highest score among several identifications derived from the respective peptide. The peptide with Cys residue at the N-terminus was excluded from the analysis due to the difficulty distinguishing the modification at the amide group or thiol group.

The XIC area values were calculated as described previously^4^. The relative peak area is that the peak area of a peptide in each concentration is divided from the highest area of the respective peptide.

## Results and Discussions

### The number of 2-ME-adducted peptides under varying concentration of 2-ME

In the absence of 2-ME, surprisingly, although the intensities of the modified Cys-containing peptides were quantitatively undetected or quite lower, the modified Cys residue in 6 and 10 peptides were identified with the molecular mass of + 76 (Fig. 1A). β-Gal samples derived from two different lots were digested without any additives and measured by MS, and their peptides modified with 2-ME were identified in each lot. Since no disulfide bond forms in β-Gal, it is speculated that 2-ME was added during the manufacturing process to prevent the oxidation of Cys residues (Supplementary Table 1).

**Figure 1.**
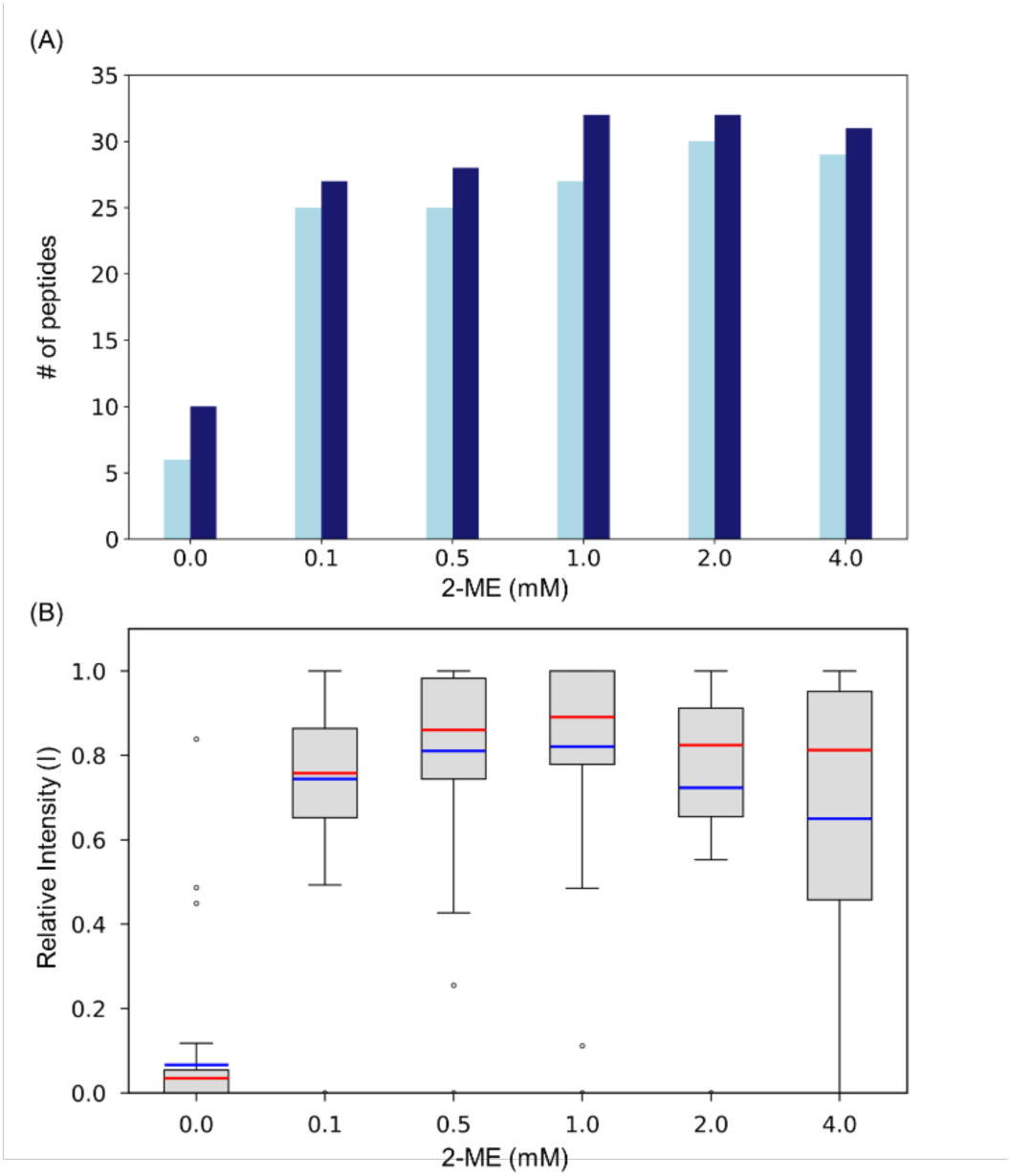
Evaluation of 2-ME adduction at varying concentration of 2-ME. (A) The number of identified 2-ME-adducted peptides in varying concentrations of 2-ME were shown. (B) Box plots representing relative peak areas of quantified 2-ME-adducted peptides at varying concentrations of 2-ME. The peptides with peak areas >10^8^ were used in this plot. The box was indicated the interquartile range (IQR) in between 25 and 75 percentiles, respectively. The median and mean values were depicted as red and blue lines, respectively. The minima and maxima within 1.5 times the IQR below 25 and above 75 percentiles were exhibited as whiskers, respectively. Outliers, signified by individual dots, fall outside the bounds.

The efficiency of 2-ME adduction to the Cys residue varying concentration of 2-ME was evaluated. Increasing the concentration of 2-ME was provided increasing number of 2-ME-adducted peptides and then the number of the peptides reached the plateau with around 30 peptides at the concentration of 2-ME higher than 0.5 mM, suggesting that the addition of 0.5 mM 2-ME was enough to modify the Cys residue efficiently (Fig. 1B).

### Assessment of DMSO concentration for 2-ME adduction

To confirm the DMSO promoted the adduction of 2-ME to the Cys residue. Figure 2A shows the number of peptides modified by 1 mM 2-ME under varying concentrations of DMSO and of carbamidomethylated peptides using IAA, respectively. 2-ME adduction was identified under all conditions even without the DMSO. 48 peptides were identified as 2-ME adduction at 0% DMSO. Increasing the concentration of DMSO affected the increasing number of the peptide with 2-ME adduction, and thus the number of the peptide with 2-ME adduction reached 132 at 30% DMSO, suggesting that DMSO promoted the 2-ME adduction for the thiol group in the Cys residue depending on the concentration of DMSO. 982 peptides without Cys were also assigned at 0% DMSO, but the increase in the concentration of DMSO decreased the number of the identified peptides to 841 at 30% DMSO. In contrast to the number of peptides with 2-ME adduction, the number of the peptides with Free Cys residue was decreased depending on the increase of the DMSO concentration, subsequently, no peptide with Free Cys was observed at 30% DMSO.

**Figure 2.**
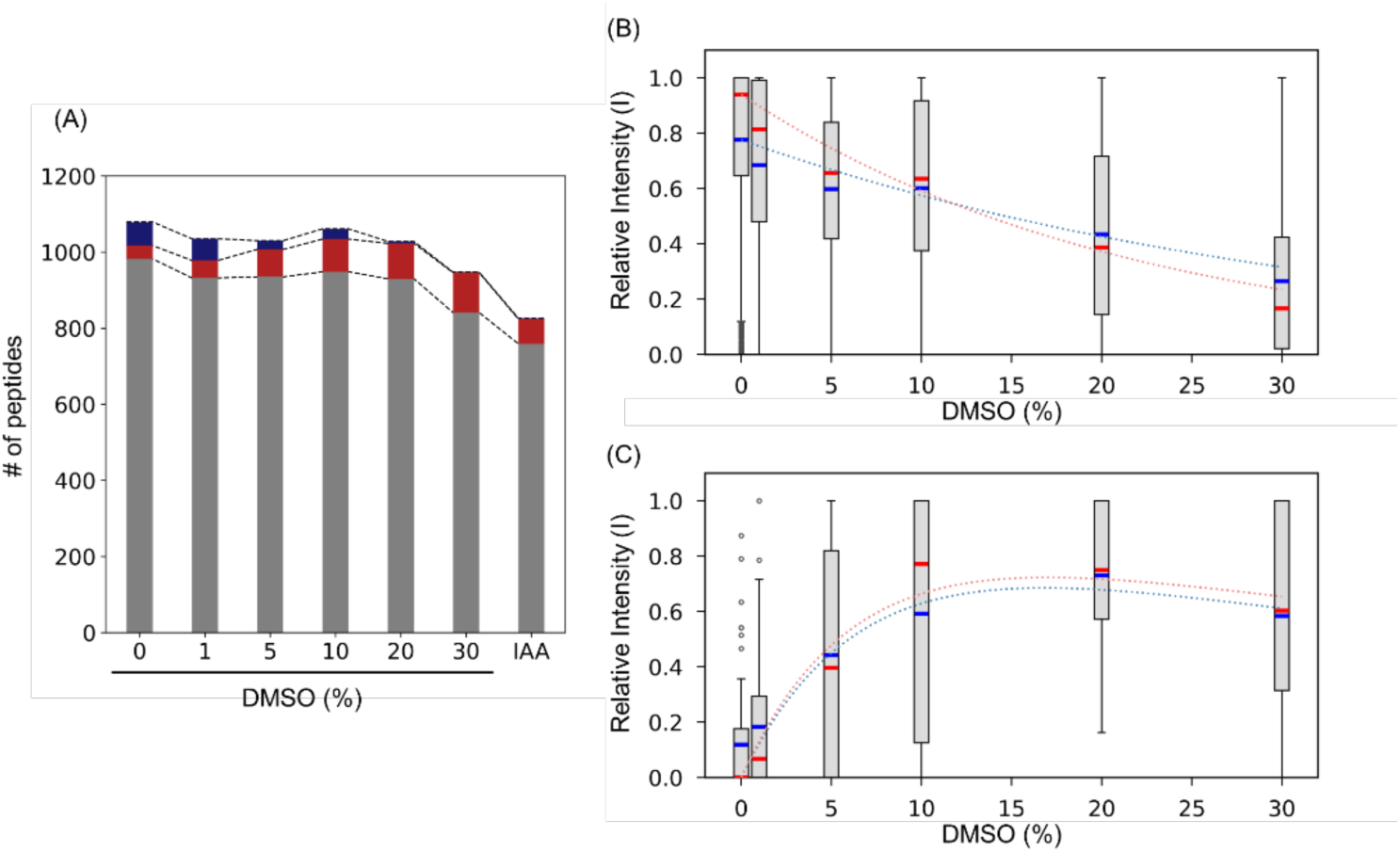
Efficiency of 2-ME adduction at varying concentration of DMSO. (A) The number of identified peptides at varying concentrations of DMSO or IAA were shown. The peptides without Cys, modified Cys, and non-modified Cys were depicted as gray, red, and blue bars, respectively. Box plots representing relative peak areas of (B) without Cys at varying concentrations of DMSO and (C) quantified peptides with Cys adducted by 2-ME, respectively. The peptides with peak areas >10^8^ were used in this plot. This figure is displayed in the same manner as in Fig. 1.

On the other hand, 844 peptides were identified under the IAA treatment commonly used for the alkylation of Cys for proteomics. However, the number of the identified peptides treated with IAA was 10% lower than those of identified peptides at 30% DMSO.

### Relative peak areas of 2-ME-adducted β-Gal at varying DMSO concentrations

The number of 2-ME-adducted peptides was found to increase in a DMSO concentration-dependent manner. The trend in relative peak areas of identified 2-ME-adducted peptides with varying concentrations of DMSO was also evaluated. The relative peak areas of peptides without Cys at 0% DMSO indicated mean and median values of 0.78 and 0.94, respectively (Fig. 2B). Increasing the concentration of DMSO up to 30%, the mean and median values dropped down to 0.27 and 0.17, respectively, decreasing the intensity exponentially as *e*^(−0.03[DMSO])^ in the average and *e*^(−0.05[DMSO])^ in the median, respectively. The previous investigation exhibited that DMSO reduced the optimal temperature and promoted the catalytic activity at 25°C of the chymotrypsin-like protease (3CLpro) of SARS-CoV-2^18^. In contrast to 3CLpro, the trend of the peak intensity suggested that the chymotryptic activity was suppressed in the DMSO concentration-dependent manner at 25℃. The trend in number of identified peptides was also the DMSO concentration-dependent manner, supporting the suppression of the chymotryptic activity depending on the concentration of DMSO. In the absence of DMSO, 35 peptides adducted with 2-ME were identified but their relative peak areas were quite small or hardly detected with mean and median values of 0.12 and 0, respectively (Fig. 2C), suggesting that the 2-ME adduction was not catalyzed in the absence of DMSO.

Increasing the concentration of DMSO promoted the 2-ME adduction, and the relative peak areas of the 2-ME-adducted peptides reached mean and median values of 0.73 and 0.75 at the 20% DMSO, respectively, allowing the quantification of 72 2-ME-adducted peptides with intensities >10^8^. Increasing the concentration of DMSO up to 30% decreased the relative peak areas of the 2-ME-adducted peptides with mean and median values of 0.58 and 0.60, respectively. The trend in the relative peak areas of 2-ME-adducted peptides depending the concentration of DMSO exhibited exponentially as {1 – *e*^(−0.13[*DMSO*])^}*e*^(−0.02[*DMSO*])^ for the mean value, indicating the product of a term showing a concentration-dependent increase in area values, {1 – *e*^(−0.13[*DMSO*])^}, and a term showing a concentration-dependent decrease in area values, *e*^(−0.02[*DMSO*])^. The rate of decrease in relative peak areas of 2-ME-adducted peptides containing Cys, *e*^(−0.02[*DMSO*])^, was similar to that for peptides without Cys, *e*^(−0.03[*DMSO*])^, suggesting that the decrease rate may indicate the chymotryptic activity depending on the concentration of DMSO and the increasing rate may exhibit the modification rate of 2-ME adduction by DMSO. The trend of median value was also similar manner to the average one and was represent as {1 – *e*^(−0.14[*DMSO*])^}*e*^(−0.01[*DMSO*])^.

### Confirmation of offsite modification in 2-ME adduction

Under the IAA treatment, over 25 peptides were identified as the carbamidomethylation to Cys residue in two independent experiments (Fig. 3). The carbamidomethylation at the N-terminus was also observed with over 100 peptides, as more pronounced modifications rather than for Cys residue, and thus the N-terminus was found to be the most carbamidomethylated. Furthermore, the software detected more than 10 peptides as modifications to the other amino acids; histidine (His), Met, aspartic acid (Asp), glutamine (Gln), arginine (Arg), Tryptophan (Trp), asparagine (Asn), and threonine (Thr). The carbamidomethylation for His, Asp, and Thr residues is known^7^ and Met residue is carbamidomethylated at lower pH^8^. However, the *m*/*z* values of Gln, Trp, Arg, and Asn were also increased by +57 Da in two independent experiments, suggesting that they suspected to be carbamidomethylated.

**Figure 3.**
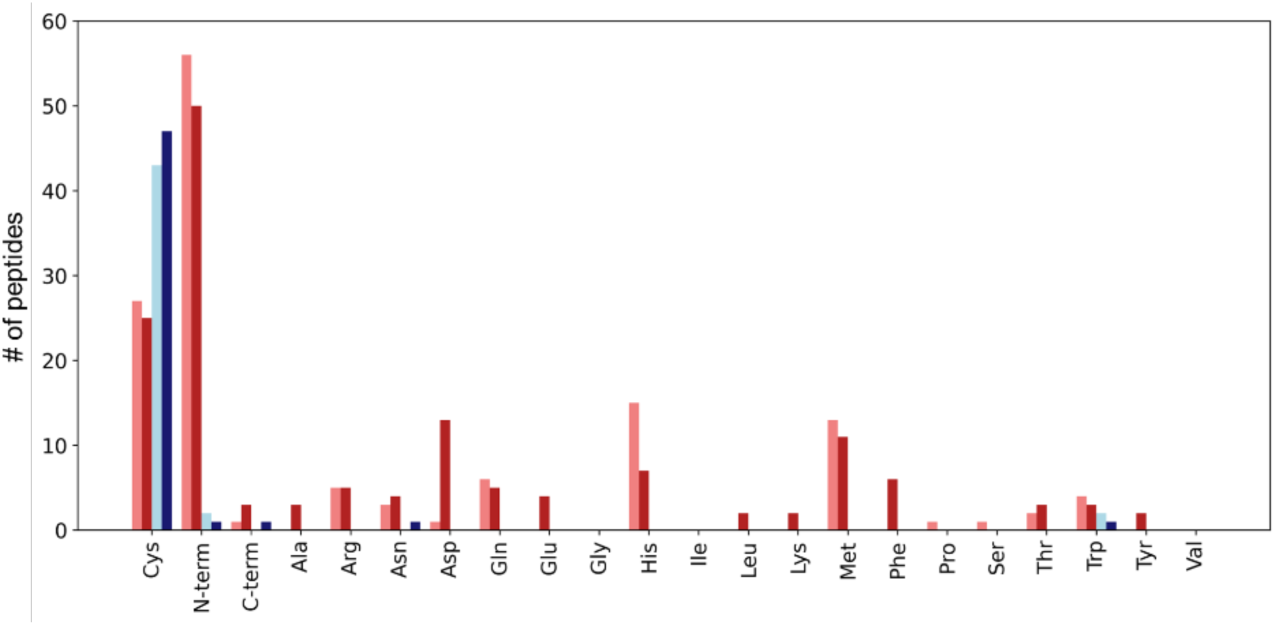
The side reactions compared between 2-ME adduction and IAA. The number of the peptides modified at each amino acids in the presence of 20% DMSO and IAA were displayed, respectively. Two independent experiments both DMSO and IAA treatments were performed. The number of peptides containing modification using DMSO while the duplicate experiments were depicted as sky blue and blue bars, and those treated with IAA were indicated as salmon and red, respectively.

On the other hand, in the 2-ME catalytic reaction promoted by DMSO, a total of 90 peptides including 2-ME adducted Cys residue was observed in two independent experiments. A few peptides also obtained the increase of +76 Da at other residues by the PEAKS10 search in two independent experiments; three peptides at N-terminus modification, one peptide at C-terminus modification, and three and one peptides modified at Trp and Asn residue, respectively. Although the peptides, NRLTDDPRWLPAMSERVTRM and AHAMGNSLGGFAKYW, don’t contain Cys residue, an increase in +76 Da at Trp residue was observed in the MS/MS spectra (Supplementary Fig. 1). In contrast to these peptides, other peptides, which increased by +76 Da causing the 2-ME adduction, includes Cys residue but the modification was assigned at the other amino acid instead of Cys. Furthermore, the modifications to the residue at N-terminus, C-terminus, and Asn in these peptides were doubted, because uncertain y- and b-series were obtained (Supplementary Fig. 2). The MS/MS spectra derived from the same peptides were carefully assessed, consequently one of the MS/MS spectra clearly mentioned that Cys residue in the peptides was increased in its mass with +76 Da and thus the modification to Cys residue was assigned well rather than that for the N-terminus, C-terminus, Asn, and other Trp residues identified poorly by their MS/MS spectra. Therefore, we concluded that the modifications to the N-terminus, C-terminus, and Asn were misidentified and that 2-ME actually modified the peptides for the Cys residue.

## Conclusions

The thiol group of Cys undergoes various modifications, such as disulfide bond, sulfinylation, and sulfonylation, and thus causes complicating the MS spectra. To prevent the complicated modification during experiment, the disulfide bond is cleaved by reducing agent, 2-ME, dithiothreitol, and TCEP, subsequently the thiol group is protected using alkylating reagent before LC-MS analysis. However, alkylating reagent commonly used in proteomics, such as IAA, modifies not only Cys residue but also other amino acids and N-terminus promiscuously, causing the complicated MS spectra and then the result of the sequence search and quantitative peptide analysis was worth owing to decreasing the intensities of the MS spectra. The previous investigation in the organic chemistry suggested that DMSO promote and enhanced the oxidation of the thiol group to form such as a disulfide bond^11^. In this study, to overcome the promiscuous modification problem, we adopt a Cys-specific modification method based on this phenomenon for the protecting the thiol group in the protein. The specificity and the efficiency of the 2-ME modification by DMSO was assessed, resulting in almost all Cys residues were protected by 2-ME efficiently under 20% DMSO and 0.5 mM 2-ME. Furthermore, this method prevented the promiscuous modification and achieved the quite high specificity compared to traditional methods, making it a promising new approach for Cys modification. This technique is expected to improve the quantitative performance of protein analysis by LC-MS.

## Author contributions

T.M. conceptualized this work. A.S. performed the experiment. A.S. Y. K. and T.M. analyzed the data, wrote the paper, and reviewed the results and approved the final version of the manuscript.

## Acknowledgment

This work was supported by Grants-in-Aid for Scientific Research from the Ministry of Education, Culture, Sports, Science and Technology, Japan (grant numbers 17KK0141, 21K06036), AMED, and NEDO.

## Conflict of Interest

The authors declare that they have no competing financial interests.

## Notes

### Competing Interest Statement

The authors have declared no competing interest.

