## Supplementary Figure 1 for "Reducing Offsite Modification using 2-mercaptoethanol for Proteome Analysis"

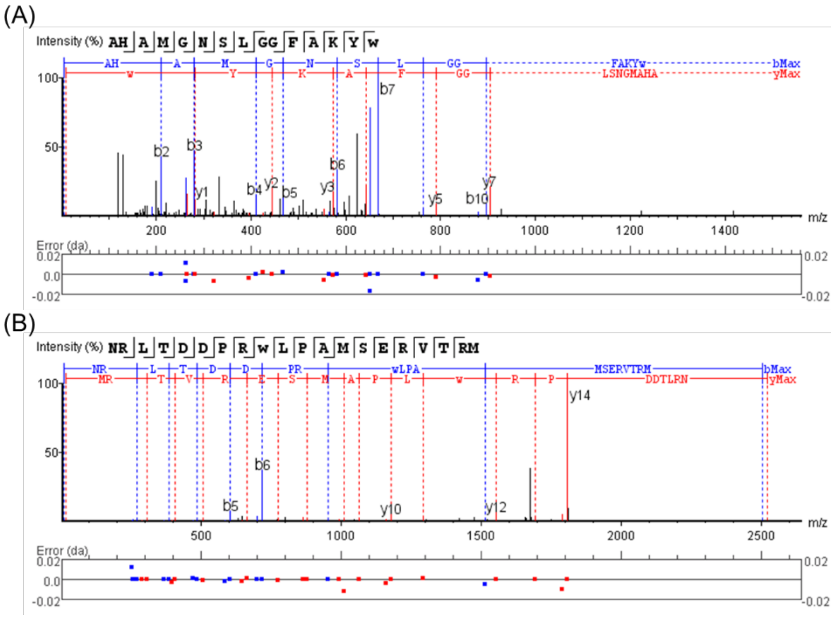


**Supplementary Figure 1**. MS/MS spectra of a peptide containing modified Trp residue speculated as side reaction. MS/MS spectra with sequence assignments of fragment ions corresponding to (A) "AHAMGNSLGGFAKYW(+76)" with an *m*/*z* 562.5905 (*z* = 3) and (B) "NRLTDDPRW(+76)LPAMSERVTRM" with an *m*/*z* 631.0629 (*z* = 4), respectively. MS/MS spectra were deconvoluted into singly charged ions from the observed spectra and peaks were assigned theoretical *m*/*z* values for fragment ions. The annotations of the identified matched N-terminal-containing ions are shown in blue and the C-terminal-containing ions in red, respectively. The *m*/*z* differences between theoretical and observed values for all assigned peaks were displayed in error map and were less than 0.02 Da.
