## Supplementary Figure 2 for "Reducing Offsite Modification using 2-mercaptoethanol for Proteome Analysis"

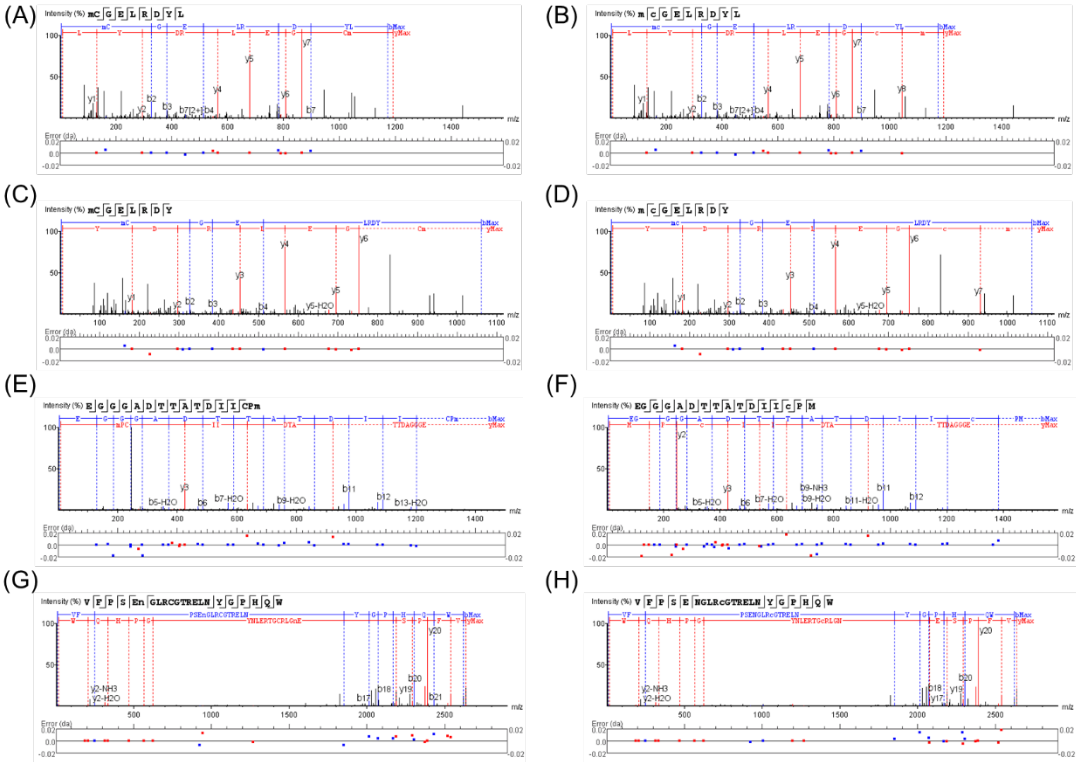

**Supplementary Figure 2**. MS/MS spectra of a peptide containing Cys residue identified as a side reaction. MS/MS spectra with sequence assignments of fragment ions corresponding to (A) “m(+15)(+76)CGELRDYL” with an *m*/*z* 596.2461 (*z* = 2), (B) “m(+15)c(+76)GELRDYL” with an *m*/*z* 596.2461 (*z* = 2), (C) “m(+15)(+76)CGELRDY” with an *m*/*z* 539.704 (*z* = 2), (D) “m(+15)c(+76)GELRDY” with an *m*/*z* 539.704 (*z* = 2), (E) “EGGGADTTATDIICPm(+76)” with an *m*/*z* 814.3358 (*z* = 2), (F) “EGGGADTTATDIIc(+76)PM” with an *m*/*z* 814.3381 (*z* = 2), (G) “VFPSEn(+76)GLRCGTRELNYGPHQW” with an *m*/*z* 879.4086 (*z* = 3), (H) “VFPSENGLRc(+76)GTRELNYGPHQW” with an *m*/*z* 879.4086 (*z* = 3). This figure is displayed in the same manner as in Supplementary Figure 1.
