## Supplementary Table 1 for "Reducing Offsite Modification using 2-mercaptoethanol for Proteome Analysis"

**Supplementary table 1**. The total number of peptides measured in two independent measurements of β-Gal from different lots without 2-ME and/or DMSO treatment. 2-ME-adducted peptides were identified in both measurements.

| | |
|---|---|
| No. of Peptide without Cys | 668 |
| No. of 2-ME-adducted peptide | 5 |
| No. of Non-modified Cys peptide | 17 |
